# POSSIBLE MULTIPLE ORIGINS OF SOME IMPORTANT CHARACTERISTICS OF THE KEEL (PAPILIONATE) FLOWERS WITHIN FABALES

**DOI:** 10.1101/2024.10.29.621004

**Authors:** Deniz Aygören Uluer

## Abstract

Keel flowers are bilaterally symmetrical, pentamerous flowers with the reproductive organs enclosed by keel petals. Within Fabales, keel flowers are dominant in two species-rich lineages, tribe Polygaleae (Polygalaceae) and subfamily Fabaceae (Papilionoideae); however, independent events are also observed, such as in the genus *Cercis*. Before phylogenetic advancements were available (i.e., in contrast to more recent studies), most of the studies hypothesized a non-keeled origin for the Faboideae, although a detailed investigation has never been carried out. In this study, using the results of Aygören Uluer *et al*. (2020, 2022a), the origin of some important morphological characters of the keel flower are examined, namely floral symmetry, perianth heteromorphism (i.e., three distinct petal/+sepal types), and the presence of enclosed reproductive organs. These characters are analysed within the Fabales using three different ancestral state analyses based on a phylogeny constructed for 678 taxa using published *matK*, *rbcL* and *trnL* plastid gene regions. The analyses show that symmetry probably originated in the (Fabaceae+Polygalaceae) clade, while the enclosed reproductive organs and three-types of petals appear to have evolved independently multiple times. Interestingly, not only the enclosed reproductive organs but also petal heteromorphism probably did not evolve in the MRCA of the Faboideae, but rather in a very early stage of the evolution of the subfamily. While future homology assessments and/or evolutionary developmental genetic (evo-devo) studies will be required to more clearly elucidate the evolutionary processes, the current study is the first attempt to investigate the origin of some important characteristics of keel flowers within order Fabales.

## INTRODUCTION

Keel flowers (*sensu* Westerkamp, 1997) are (mostly) zygomorphic, pentamerous flowers with three different petal types (a standard or vexillum, two wings or alae and a keel or carina comprising two usually partially fused petals), with the reproductive organs enclosed within the protective keel (Polhill & Raven, 1981; Endress, 1996; Pennington *et al*. 2000; Persson, 2001; Tucker, 2002; Citerne *et al*. 2006; Westerkamp & Claßen-Bockhoff, 2007; Carvalho *et al*. 2023). Within the Fabales Bromhead, unlike the actinomorphic flowers of the species-poor families Surianaceae Arn. and Quillajaceae D. Don, keel flowers are dominant in two species-rich lineages: tribe Polygaleae Chodat (Polygalaceae Hoffmanns. & Link) with ca. 800 species and subfamily Faboideae DC. (Fabaceae Lindl.) with ca. 14,000 species (Lewis *et al*. 2005; Bello *et al*. 2007, 2010; LPWG, 2017). While keel flowers are also found outside of Faboideae and Polygaleae, such as in the legume subfamilies Cercidoideae LPWG, Dialioideae LPWG and Caesalpinioideae DC. and in many unrelated families, including Ranunculaceae Juss., Onagraceae Juss., Sapindaceae Juss, Trigoniaceae A. Juss., Geraniaceae Juss., Tropaeolaceae Juss. ex DC., Papaveraceae Juss., Solanaceae Juss., Acanthaceae Juss. and Commelinaceae Mirb. (with the exception of Trigoniaceae and Fumarioideae, keel flowers are not as common as in Fabales (Arroyo, 1981; Polhill *et al*. 1981; Westerkamp, 1997; Westerkamp & Weber, 1999).

The 16 independent origins of keel flowers within 10 different angiosperms orders, both in monocots and eudicots (Westerkamp, 1997), has been referred to as an adaptive response to bees (Leppik, 1966; Faegri & Van der Pijl, 1979; Arroyo 1981; Westerkamp, 1997; Westerkamp and Weber, 1999; Córdoba & Cocucci 2011; Etcheverry *et al*. 2003; Amaral-Neto *et al*. 2015). The keel flowers evolved to attract bees as legitimate pollinators and to protect the pollen, from robbery by undesirable flower visitors, by forming a closed envelope around the stamens (Arroyo, 1981; Westerkamp, 1996, 1997; Westerkamp & Weber, 1999). While keel flowers are widely distributed across many angiosperm orders (Westerkamp, 1997), due to the developmental differences and different evolutionary modules of the keeled flowers in different clades (Bello *et al*. 2010, 2012), this floral trait has been recognised as an adaptation or convergent character rather than a synapomorphy (Westerkamp, 1997; Prenner, 2004; Bello *et al*. 2010). For instance, in both subfamily Faboideae and tribe Polygaleae, keel flowers are 5-merous, and consist of three parts: a standard (banner or vexillum) petal which is composed of single petal in Fabaceae but replaced by sepals in Polygalaceae, two wing petals (alae) in Faboideae replaced by two petaloid sepals in Polygalaceae, and two keel petals (together forming the carina) which are partially fused in Faboideae, but represented by a single petal in Polygalaceae (Arroyo, 1981; Eriksen, 1993; Westerkamp, 1997; Westerkamp and Weber, 1999; Persson, 2001; Tucker, 2003a; Prenner, 2004). A legume exception with a papilionate flower is the cercidoid genus *Cercis* L. (Figure 1N). Polhill *et al*. (1981) used the term “pseudo-papilionoid” for *Cercis* flowers which resemble the Pailionoideae keel flowers, they are both bilateraly symmetrical, have three different petal types, and enclosed reproductive organs; although *Cercis* flowers lack fused stamens and a tripping mechanism (Tucker, 2002) and the standard petal is innermost rather than outermost. In addition to tribe Polygaleae, keel flowers have been reported in other taxa of the Polygalaceae. For instance, Van der Meijden (1982) reported that some *Xanthophyllum* Chodat (Figure 1O) species can have keel flowers. Likewise, Breteler and Smissaert Houwing (1977) reported that *Carpolobia* G. Don (Figure 1P) and *Atroxima* Stapf flowers display bilateral symmetry and a “keel petal” which encloses the style and the stamen sheath, similar to those of Faboideae keel flowers.

**Figure 1.**
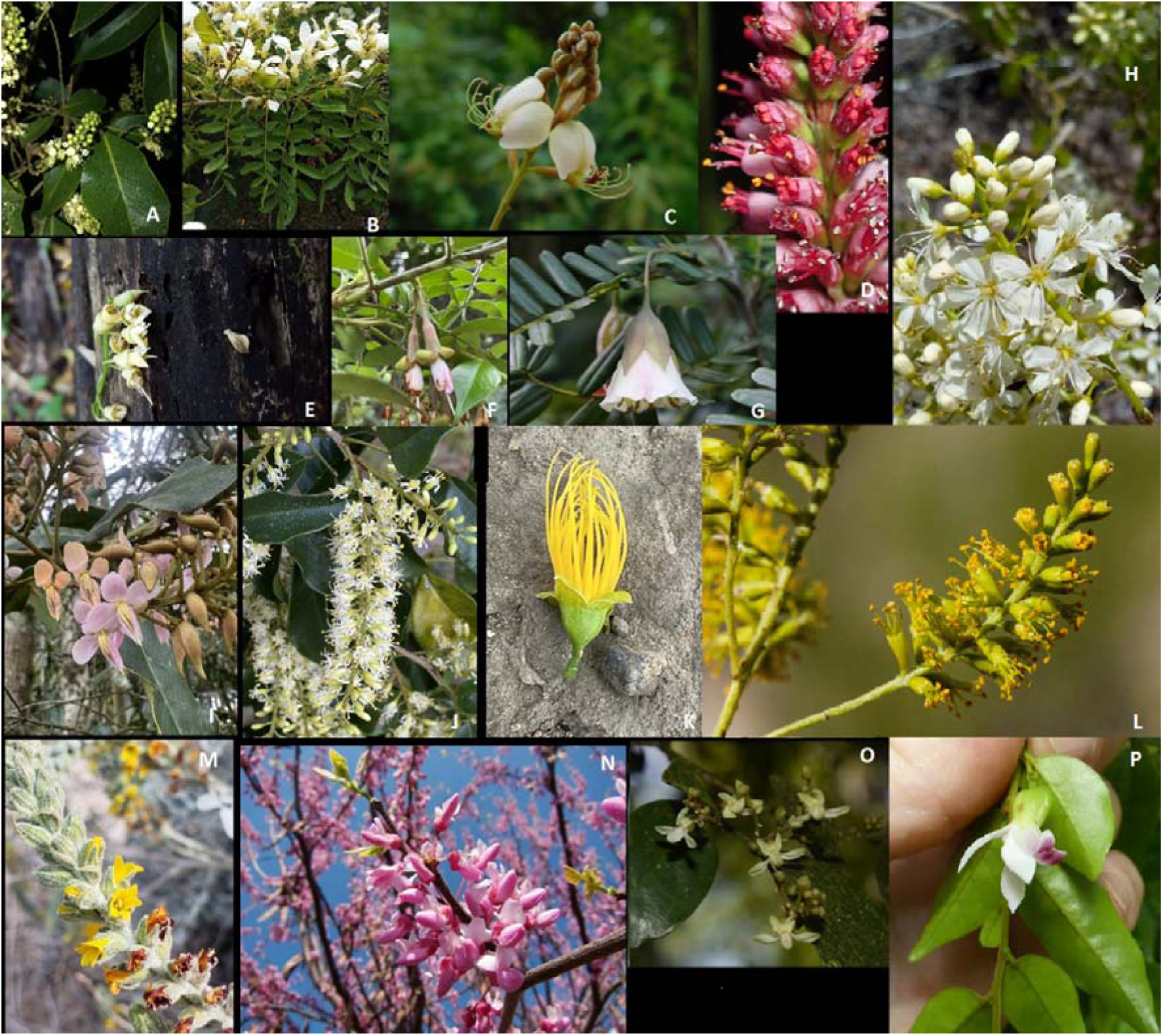
Examples of various plants that illustrate key characteristics are discussed in this study. A*- Ateleia* (Image by leo_rquiros, https://inaturalist.ca/observations/101360705), B-*Cyathostegia* (Image by Bodo Nuñez Oberg, https://inaturalist.ca/photos/31331115), C-*Amburana* (Image by Ignacio Barrientos, https://inaturalist.ca/observations/101986718), D-*Amorpha* (Image by Jared Shorma, https://inaturalist.ca/observations/168533567), E- *Lecointea* (Image by sarievanbelle, https://inaturalist.ca/observations/37404460), F-*Exostyles* (Image by Geovane Siqueira, https://inaturalist.ca/observations/168971786), G- *Cadia* (Image by Stuart Cable, at https://inaturalist.ca/observations/827974), H- *Dicraeopetalum*(Image by Solofo Eric Rakotoarisoa, https://inaturalist.ca/observations/69252906), I-*Zollernia* (Image by Funez, https://inaturalist.ca/observations/243354938), J- *Myrocarpus* (Image by Geovane Siqueira, https://inaturalist.ca/observations/225697338), K- *Cordyla* (Image by Ingolf Askevol, https://inaturalist.ca/observations/186531954), L- *Parryella* (Image by Cecelia Alexander, https://inaturalist.ca/observations/50100832), M- *Errazurizia* (Image by Joey Santore, https://inaturalist.ca/photos/108196373), N-*Cercis* (Image by Shihmei Barger, https://inaturalist.ca/photos/20246), O- *Xanthophyllum*(Image by Siddarth Machado, https://inaturalist.ca/photos/66230653), P- *Carpolobia* (Image by Carel Jongkind, https://inaturalist.ca/observations/201278807).

Subfamily Faboideae is largely characterized by keel flowers (LPWG, 2017). Indeed, the subfamily mainly differs from the other five Fabaceae subfamilies in having this flower type. In addition, in the papilionoid keel flower the standard is the outermost petal while it is innermost in subfamily Caesalpinioideae and better referred to as the median petal. Most papilionoid flowers have their sepals united into a calyx tube (LPWG, 2017). However, some early branching lineages (note that this term is used here as a working hypothesis, not for the depiction of evolution), including Sophoreae (Spreng. ex DC. 1825) Cardoso *et al*. 2013, Swartzieae DC., Dipterygeae Polhill and some Dalbergieae Bron ex DC. (Cardoso *et al*. 2013) have most, or many non-keeled flowers with radial symmetry, free stamens and unspecialized petals and together display more floral diversity than most other Faboideae (Polhill, 1981; Pennington *et al*. 2000, 2001; Lavin *et al*. 2001; Doyle and Luckow, 2003; Tucker, 2003b; Cardoso *et al*. 2012a; Klitgård *et al*. 2013; Zimmerman *et al*. 2017). For example, with respect to petal number, *Baphiopsis* Benth. ex Baker flowers have six petals, *Aldina* Endl. 3–6 petals, *Holocalyx* Micheli 5–6 petals, while *Ateleia* (DC.) Benth. (Figure 1A), *Cyathostegia* (Benth.) Schery (Figure 1B), *Amburana* Schwacke & Taubert (Figure 1C), and *Amorpha* L. (Figure 1D) all possess only one petal. With regard to stamen number, *Ateleia* flowers have 6-30 stamens, *Cyathostegia* 20-30 stamens, and *Holocalyx* 12–16 stamens. Considering petal differentiation and symmetry, *Lecointea* Ducke (Figure 1E), *Zollernia Wied-Neuw. & Nees*, *Harleyodendron* R.S. Cowan, *Exostyles* Schott (Figure 1F), and *Holocalyx* generally have flowers with five un-differentiated petals, *Cadia* Forssk. (Figure 1G), *Amphimas* Pierre ex Harms, *Acosmium* Schott, *Myrocarpus* Allemão, *Holocalyx, Riedeliella* Harms and *Dicraeopetalum* Harms (Figure 1H) have undifferentiated/poorly differentiated petals with radial/ slightly bilateral symmetry, while *Cordyla* Lour. (Figure 1K)*, Parryella* Torr. & A. Gray (Figure 1L)*, Errazurizia* Phil. (Figure 1M) have only two petal typesand *Mildbraediodendron* Harms. has no petals (Pennington *et al*. 2000; McMahon and Hufford, 2005). Furthermore, ontogenetic studies have shown that while flowers of *Zollernia* (Figure 1I), *Myrocarpus* (Figure 1J), *Lecointea*, *Harleyodendron*and *Exostyles* have five un-differentiated petals and ten stamens, these arise from fused tissue that surrounds the ovary, while in *Amburana* flowers one petal and ten stamens arise from a deep hypanthium (Pennington *et al*. 2000).

Due to the strong resemblance in flowers of some of these above-mentioned papilionoid genera to Caesalpinioideae flowers, the tribes Sophoreae and Swartzieae, as traditionally circumscribed, were considered to be “transitional groups” between Caesalpinioideae and Faboideae (Polhill *et al*. 1981; Ireland *et al*. 2000; Pennington *et al*. 2001). Whilst tribe Sophoreae was considered as a “tribe of convenience” (Polhill, 1981), tribe Swartzieae was later referred as “transitional” between Caesalpinioideae and Faboideae (Polhill, 1994). However, more recent studies have revealed that, apart from tribe Dipterygeae (part of the ADA clade of Cardoso *et al*. 2012a (i.e., the Angylocalyx Taub., Dipterygeae, Amburana clade) and Swartzieae sensu stricto, these previously recognised supra-generic taxa are not monophyletic, but rather are scattered across the Faboideae phylogenetic tree (Tucker, 1994; Doyle *et al*. 1997; Ireland *et al*. 2000; Pennington *et al*. 2000, 2001; Tucker, 2002; Cardoso *et al*. 2012a; Cardoso *et al*. 2013; Choi *et al*. 2022). As well as in some early branching papilionoid lineages, non-keeled flowers are also found in tribe Amorpheae (McMahon and Hufford, 2004), some Dalbergioids (Lavin *et al*. 2001; Cardoso *et al*. 2012b), Genistoids (Cardoso *et al*. 2012a), Lecointeoids (Mansano *et al*. 2004), and Vataireoids (Cardoso *et al*. 2013).

Until phylogenetic analyses, most studies, based largely on morphology, e.g., Arroyo (1981) and Polhill & Raven (1981), hypothesized a non-keeled origin for the papilionoid flower, in contrast to more recent studies (e.g., Pennington *et al*. 2000; Lavin *et al*. 2001; Citerne *et al*.2006; Cardoso *et al*. 2012a; Carvalho *et al*. 2023). Pennington *et al*. (2000) suggested that twelve independent reversals from the keel flower type to a non-keel type occured within the evolution of the Faboideae. Nevertheless, the ancestral floral type of Faboideae is still uncertain (Leppik, 1966; Pennington *et al*. 2000, 2001; Lavin *et al*. 2001; Wojciechowski, 2003; McMahon & Hufford, 2004; Cardoso *et al*. 2013; Klitgård *et al*. 2013; Amaral-Neto *et al*. 2015). Similarly, a comprehensive investigation into the origin of keel flowers within the Polygalaceae has not been undertaken. Several questions remain unanswered, including how many times, and when, keeled and non-keeled flowers evolved within the order Fabales? Was the origin of flower type in subfamily Faboideae and family Polygalaceae keeled or non-keeled? Do the the ancestral keel flowers resemble the extant ones? How many flower type character state reversals have there been? Which ecological and/or evolutionary factors triggered the evolution of keel flowers, and when did these events take place?

A recent study by Aygören Uluer *et al*. (2022a) conducted extensive ancestral character and ancestral area analyses to identify a possible mimicry scenario between the legume subfamily Faboideae and family Polygalaceae. The study primarily focusses six Faboideae clades, rather on the whole order Fabales. Furthermore, the authors exclusively employed MCMC ancestral state analyses and focused on those morphological characters that are visually evident to pollinators (e.g., plant height, inflorescence size, floral size), while neglecting characters important for the definition of keel flowers. Consequently, the study failed to address the possible origins of key aspects of keel and/or non-keeled flower evolution within the Fabales.

In the literature, three distinct approaches have been used to address flower evolution within lineages: the examination of the fossil record, evolutionary developmental genetic (evo-devo) studies, and the reconstruction of ancestral states utilizing extensive datasets (Sauquet *et al*. 2017). Here, I opted for the latter approach—a detailed exploration conducted through three distinct ancestral character reconstructions focusing on three morphological traits, complemented by molecular dating analyses.

## MATERIAL AND METHODS

### Taxon sampling, Alignment and Phylogenetic Analyses

I used the *mat*K, *trn*L and *rbc*L plastid matrix for ancestral trait analysis, and the results of a Maximum Likelihood (ML) and molecular dating analyses of Aygören Uluer *et al*. (2020a, 2022a). The matrix includes three plastid gene regions for 615 Fabaceae species, 14 Polygalaceae species, five Surianaceae and the sole genus of Quillajaceae, *Quillaja* (total 678 taxa. I sampled ca. 80% of Fabaceae genera (ca. 3% of the total species number) and 70% of Polygalaceae genera (1.4% of the total species number).

I accepted the phylogenetic classifications of Lewis *et al*. (2005), Gagnon *et al*. (2016) and LPWG (2017). The monophylly of the order Fabales and family Fabaceae have been strongly supported (e.g., Forest *et al*. 2007; Bello *et al*. 2009, 2012; Aygören Uluer *et al*. 2020a; 2020b; Koenen *et al*. 2020, 2021); however, the internal phylogenetic relationships are not stable, both the topology and the root change depending on the choice of genes, outgroups and methods; this possibly due to rapid radiation (Aygören Uluer *et al*. 2020a). To account for the unresolved phylogenetic relationships within Fabales, four ancestral state reconstructions I used the sample of bootstrap trees with branch lengths.

For sequence alignment and trimming I used Geneious Pro 4.8.4 (Kearse *et al*. 2012). All indels were treated as missing data. By using jModelTest2.1.10 (Guindon and Gascuel, 2003; Darriba *et al*. 2012) the most appropriate model was selected as GTR+G+I, for each of the genes. Maximum likelihood (ML) analyses were implemented in RAxML (Stamatakis *et al*. 2008), by conducting 1000 bootstrap replicates under a gamma model of heterogeneity.

### Molecular dating analysis

I directly used the molecular clock analysis results of Aygören Uluer *et al*. (2022a). BEAST v.1.8.0 (Drummond and Rambaut, 2007a) was implemented for the divergence time estimates based on 30 fossil (24 ingroup and 6 outgroup) calibrations, (see Aygören Uluer *et al*. 2022a for further explanations about the fossil calibrations). Forty-three outgroup taxa from Fabidae families (i.e., Celastrales, Cucurbitales, Fagales, Malpighiales, Oxalidales, Rosales, Zygophyllales) were also included. The reason for the large outgroup sampling was to obtain a more balanced tree, because sparse outgroup sampling with a dense ingroup sampling may cause over-estimation of the age of Fabales (Aygören Uluer *et al*. 2020a).

The BEAST input file was generated using BEAUti v.1.8.0 (Drummond *et al*. 2012). Calculations were performed online via the CIPRES Portal (Miller *et al*. 2010) by using the Yule process with a randomly generated starting tree and a lognormal relaxed model (Drummond *et al*. 2006). Searches were conducted with 2 × 107 MCMC generations, sampling every 1000th generation. LogCombiner v.1.8.0 (Drummond and Rambaut, 2007b) was used to compile the two independent runs and Tracer v.1.6 (Rambaut *et al*. 2014) was used to visually check for proper mixing and convergence. TreeAnnotator v.1.8.0 (Rambaut and Drummond, 2007) was used to obtain the maximum clade credibility tree.

### Ancestral Trait Analyses

Since the non-keel flowers within Faboideae are not homologous (Pennington *et al*. 2000), I focussed on three important keel flower morphological traits: perianth heteromorphism (i.e., presence of three distinct petal types and petals+sepals in Polygalaceae), presence of enclosed reproductive organs and floral symmetry (Westerkamp, 1997; Tucker, 2002; Cardoso *et al*. 2012a) (Table 1).

**Table 1:**
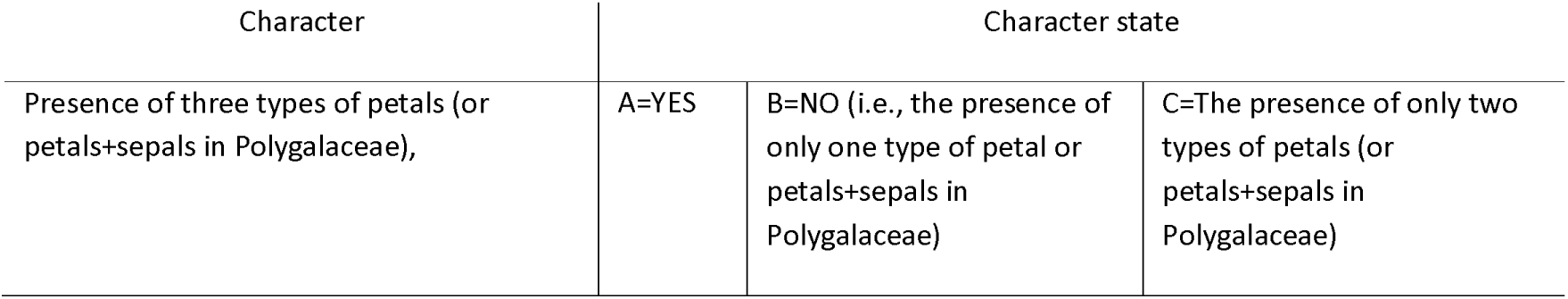
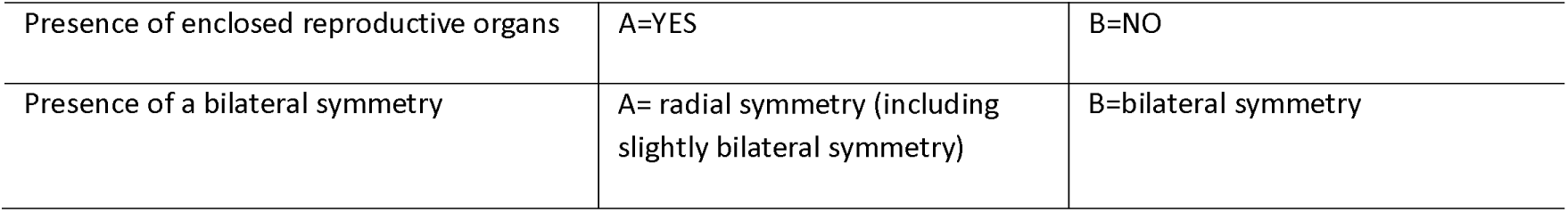
Explanation of three pollination syndrome characters coded as A, B and/or C.

| Character | Character state |  |  |
| --- | --- | --- | --- |
| Presence of three types of petals (or petals+sepals in Polygalaceae), | A=YES | B=NO (i.e., the presence of only one type of petal or petals+sepals in Polygalaceae) | C=The presence of only two types of petals (or petals+sepals in Polygalaceae) |

|  |  |  |
| --- | --- | --- |
| Presence of enclosed reproductive organs | A=YES | B=NO |
| Presence of a bilateral symmetry | A = radial symmetry (including slightly bilateral symmetry) | B=bilateral symmetry |

The definition of the papilionoid flower changes from one study to another (Tucker, 2002); however, in a broad sense having a pentamerous corolla (e.g., Tucker, 1993; Pennington *et al*. 2000; Sinjushin, 2019), petal heteromorphism (Tucker, 1993, 1997; Pennington *et al*. 2000; Prenner & Klitgård, 2008; Carvalho *et al*. 2023), presence of bilateral symmetry (Tucker, 1993, 1997; Sinjushin, 2019; Carvalho *et al*. 2023), presence of enclosed reproductive organs (Tucker, 2002, Westerkamp & Claßen-Bockhoff, 2007; Carvalho *et al*. 2023), and stamen filament connation (Tucker, 1993, Leite *et al*. 2014; 2015; Carvalho *et al*. 2023) are the most important morphological features of a keel flower. However, while I adopted three of these characters (i.e., petal heteromorphism, enclosed reproductive organs and symmetry), I did not include the other two characters as being typical characters of a keel flower, namely fused stamens and the presence of a pentamerous corolla.

Considering petal number, flowers of some taxa can have petal abortion (Tucker, 1990, 2000, 2003b; McMahon & Hufford, 2005) or petals can be initiated but later suppressed (Leite *et al*. 2015). For instance, within the Amorphoid clade, while five petals can be initiated, some are then the suppressed (e.g., in *Amorpha canescens* Pursh), or not elongated (e.g*., Amorpha fruticosa* L.), or no petal initiation is observed (e.g*., Parryella filifolia* Torr. & A. Gray) (McMahon and Hufford, 2005). On the other hand, the results of Sauquet *et al*. (2017) and the conclusion of Sinjushin (2021) confirmed that the ancestral floral form of Faboideae was pentamerous (no early-branching Faboideae lineages were included in the analyses of Sauquet *et al*. 2017). Furthermore, due to lack of homology assessments in the literature, particularly for the early branching papilonoids (Prenner *et al*. 2015) prevented me from including this character in the analyses. With regard to stamen fusion, in the early-diverging clades of Faboideae free stamens are the norm (i.e., almost all ADA and Swartzioid clade members, except four genera with fused stamens in the Dipterygeae and genus *Dussia*) (Pennington *et al*. 2000; Leite *et al*. 2015). For the fused stamens found in later evolved keel flowered lineages multiple origins have been suggested (Pennington *et al*. 2000). Moreover, the presence of fused stamens does not guarantee a keel flower formation; indeed, this character shows variation even among different keel flowered lineages, such as in the Baphieae and the Mirbelioids of the Meso-Papilionoideae (Carvalho *et al*. 2023). Therefore, I excluded these two characters from the analyses.

In addition to the characters mentioned above, many more have been attributed to keel flowers (e.g., Tucker, 1997, 2002; Leite *et al*. 2015), but the most important ones are: petal aestivation (Tucker, 2002), hypanthium formation (Tucker, 1997; Pennington *et al*. 2000), order of petal/sepal inititation (Tucker, 1997; Pennington *et al*. 2000; Leite *et al*. 2014; 2015), secondary pollen presentation (i.e., explosive, valvular, piston, brush types, Westerkamp, 1997), presence of a nectary and nectar chamber (Tucker, 1997; Westerkamp & Weber, 1999), loss of organs (Tucker, 1990), initiation and later reduction of petals (e.g., Leite *et al*. 2015), and fusion of the petals which is particularly common among later-diverging Faboideae tribes (Polhill & Raven, 1981; Tucker, 2002; Westerkamp, 1997). While some of these morphological characters are retained by only “advanced-papilionoid clades” (e.g., secondary pollen presentation, fusion among organs), others are more widely dispersed throughout the evolution of Papilionoideae (e.g., petal aestivation type, initiation and later reduction of petals). Unfortunately, it was not possible in the densely sampled taxon study to assess these characters, due to data limitation in relevant literature. The study focuses on three important characters highlighted in previous publications (i.e., the presence of three distinct petal/ petal+sepal types, the presence of enclosed reproductive organs, and floral symmetry) (Polhill & Raven, 1981; Endress, 1996; Pennington *et al*. 2000; Persson, 2001; Tucker, 2002; McMahon and Hufford, 2005; Westerkamp & Claßen-Bockhoff, 2007; Bello *et al*. 2010; Carvalho *et al*. 2023). Additionally, Van der Meijden (1982) reported that some *Xanthophyllum* Roxb. species may have keel flowers, and Breteler and Smissaert-Houwing (1977) indicated that the flowers of *Carpolobia* and *Atroxima* have a keel petal which encloses the style and the stamen sheath (all three genera in the Polygalaceae). Following Bello *et al*. (2010) and Aygören Uluer (2022a), I accept *Carpolobia* and *Atroxima* as keel flowered, but *Xanthophyllum*as polymorphic.

For the perianth heteromorphism (i.e., the presence of three types of petal, or petals+sepals in Polygalaceae) (i.e.,focussing on petal shape and size, not the function, as commonly included in the literature) analyses, I coded three states: A=YES, B=NO (i.e., the presence of only one type of petal) and C=the presence of only two types of petal (or petals+sepals in Polygalaceae). For the floral symmetry analyses, I coded two states: A= radial symmetry or slightly bilateral symmetry and B=bilateral symmetry. And for the presence of enclosed reproductive organs analyses I coded two states: A=YES and B=NO (Table 1). I directly adopted these character matrices from Aygören Uluer *et al*. (2022a) (Supporting Information, S1 to S3). In contrast to Aygören Uluer *et al*. (2022a), I employed both the population of ML and the MCMC trees. To eliminate software (RASP, BayesTraits and Mesquite) bias (i.e., conflicting results with different software), I employed three different approaches for the ancestral state analyses with populations of 100 ML trees (BayesTraits and Mesquite) or a condensed MCMC tree (RASP). These were:

1. the program BayesTraits v2.0 (Pagel & Meade, 2006) was used for Bayesian estimation of ancestral states. For the “MultiState” model, MCMC analyses were run for 2×10^6^ generations, with default settings except the rate deviation (ratedev) and RevJump (rjhp exp) parameters. Burn-in was set as the first 200,000 iterations.
2. the Bayesian binary MCMC (BBM) option (Ronquist and Huelsenbeck, 2003) of RASP v.4.2 (Reconstruct Ancestral State in Phylogenies; Yu *et al*. 2015), with default settings except the model of evolution. I ran the analyses for 1 × 10⁶generations, sampling every 100th generation, and with burn-in set to the first 2500 iterations. Due to the lilimitations of the software (i.e., taxon and tree number), I used the condensed tree which was summarized from the population of ML trees.
3. a set of analyses for the discrete characters by the unordered Maximum Parsimony (MP) option of Mesquite v3.51 (Maddison & Maddison, 2018). Interactive Tree of Life (iTOL) online tool (https://itol.embl.de/) (Letunic & Bork, 2016) was used to visualize tree file(s).

## RESULTS

### Phylogenetic Analysis

The data matrix consisted of 3,894 characters in total, and while 63% (2,445) of the characters were variable, 51% (1,968) were parsimony informative.

The ML analysis revealed that the order Fabales is monophyletic (100%), and a ((Polygalaceae (98%)+Fabaceae (99%)) (Quillajaceae+Surianaceae(98%))) topology was obtained within the order (65% and 74% BS, respectively). First, *Xanthophyllum* was sister to all remaining Polygalaceae, Carpolobieae (94%) was sister to Polygaleae (55%), and non-monophyletic Moutabeae was sister to this clade. Second, *Duparquetia* Baill. (Duparquetioideae) was sister to all Fabaceae (99% BS). Cercidoideae (99%) + Detarioideae (100%) was the second diverging clade (only 23% BS). Dialioideae (99%) was sister to remaining legumes (65%). Caesalpinioideae (90%) was sister to Faboideae (91%) with 94% BS. Within Faboideae, the ADA clade (Amburaneae+Dipterygeae+Angylocalyceae; Cardoso *et al*. 2012a) (71% BS) was sister to all remaining Faboideae, the Swartzioid clade (100% BS) was the second diverging clade, and the Cladrastis clade (96% BS) resolved as sister to the remaining Faboideae.

### Molecular dating analysis

The divergence time analysis yielded a monophyletic Fabales (1.00 PP) (Figure 2), and within the order a ((Fabaceae+Polygalaceae) (Surianaceae+Quillajaceae)) topology was generated. Within Polygalaceae, while Polygaleae and Carpolobieae were monophyletic (0.96 and 1.00 PP, respectively), tribe Moutabeae was not. Within Fabaceae, on the other hand, a (((Faboideae, 1,00 PP+Caesalpinioideae, 1.00 PP) Dialioideae, 1.00 PP) (Detarioideae, 1.00 PP (Duparquetioideae+Cercidoideae, 1.00 PP))) topology is estimated. Within Faboideae, the Swartzioid clade was sister to all remaining Faboideae (1.00 PP), the ADA clade (only 0.30 PP), was the second diverging clade.

**Figure 2.**
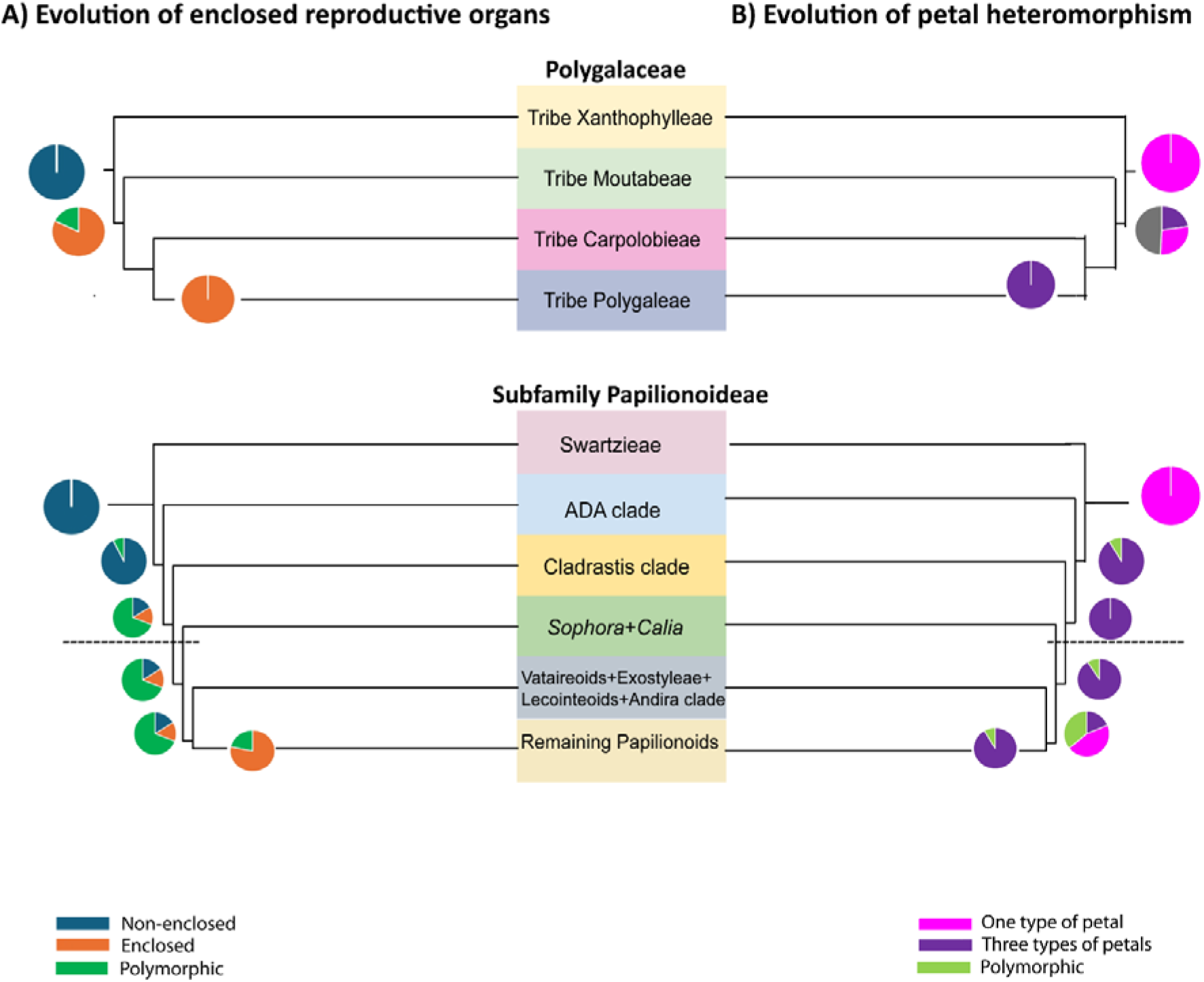
The evolution of enclosed reproductive organs (A) and petal heteromorphism (B) within the family Polylacaeae and subfamily Faboideae (Fabaceae) (note that the results of the symmetry analyses are not included due to inconclusive findings). The phylogenetic tree is pruned to show the important events within these two clades. Dashes indicate that the region (between the separation of Sophora+Calia and the remaining Faboideae) is pruned due to non-monophyly of some of the early branching lineages (note that the deleted proportions were not very different from each other). Pie charts represent the result of the RASP ancestral state analyses, while the grey section represents polymorphic+two different petal types.

The crown age of Fabales is predicted to be at least 74.97 MYA (95% HPD 69.3–76.7); the (Surianaceae+Quillajaceae) crown node as 68.62 MYA (95% HPD 50.2–73.9); the Surianaceae crown node as 47.59 MYA (95% HPD 33.2–53.1); the Polygalaceae crown node as 63.59 MYA (95% HPD 58.2– 62.7); the Polygaleae crown node as 45.16 MYA (95% HPD 38.8–44.7); the Fabaceae crown node as 71.89 MYA (95% HPD 67.9–69.3), and the Faboideae crown node as 67.19 MYA (95% HPD 62.5–64.9).

### Ancestral Trait Analyses

First, for symmetry, while the BayesTraits analysis suggested a radial or slightly bilateral symmetrical ancestor for the origin of Fabales (99.7%), the result of the RASP analysis (Supporting Information, S4) was equivocal (43.5% bilateral symmetry, and 52.5% radial or slightly bilateral symmetry) (Table 2, Figure 2). For the origin of Fabaceaee, Faboideae and family Polygalaceae while the BayesTraits (radial or slightly bilateral symmetrical origin with moderate to strong support for all nodes) and RASP BayesTraits (bilateral symmetrical origin with strong support for all nodes) analyses indicated conflicting results (Table 2). Tribe Polygaleae resolved as having a bilaterally symmetrical MRCA (Most Recent Common Ancestor) in all analyses, with moderate to strong support (79% in the BayesTraits, and 99.9% in the RASP analysis). The results of the Mesquite analyses (Supporting Information, S7), on the other hand, yielded equivocal results for the origin of Fabales, and bilaterally symmetrical ancestors for the other clades.

**Table 2.**
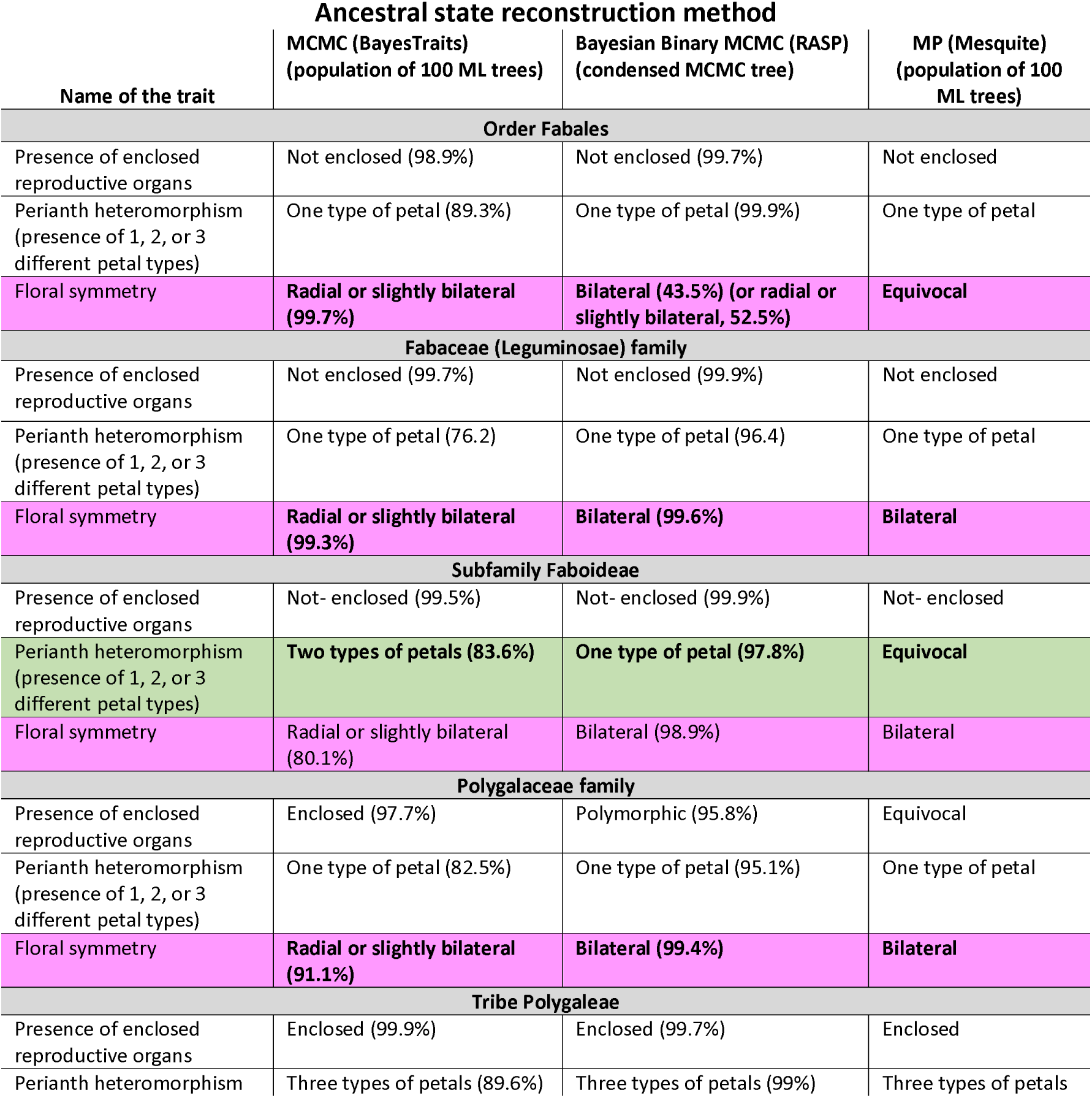
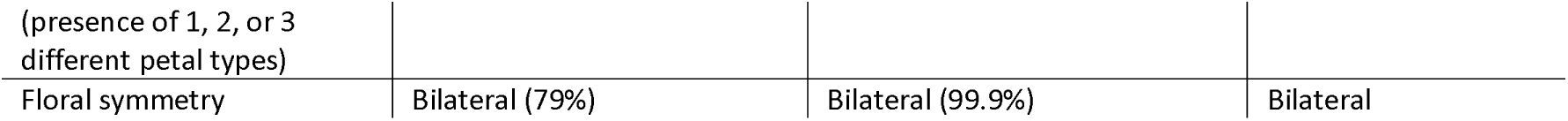
Comparison of ancestral state reconstruction analyses for the origin of Faboideae conducted with different methods. If the three analyses yielded different results, it is indicated in bold. If available, percentages are given for the results of the ancestral reconstructions.

Two RASP analyses indicated that bilateral symmetry either evolved at the origin of the order (A=52.5%, B=43.5%, node 1269) or most probably after the separation of the Quillajaceae+Surianaceae families (node 1263, A=99%), ca. 73.2 MYA, is well-preserved at the origin of Faboideae (node 1019, A=98.8%) and Polygalaceae (node 1262, A=99.4%), but lost (e.g., 13 times within Faboideae, twice within Polygalaceae and several times within the rest of the legumes) and re-gained several times within the Fabaceae+Polygalaceae clade.

Second, for the enclosed reproductive organs, all of the analyses yielded non-enclosed reproductive organ origins for order Fabales (98.9% and 99.7%, BayesTraits and RASP analyses, respectively), Fabaceae (99.7% and 99.9%), and subfamily Faboideae (99.5% and 99.9%) (Table 2, Figure 2). On the other hand, while all analyses yielded a MRCA with enclosed reproductive organs for the tribe Polygaleae (99.9% and 99.7%), for family Polygalaceae the origin was probably polymorphic (RASP analyses, node 1262, 97.4% or with enclosed reproductive organs (BayesTraits analysis, 97.7%) (note that the Mesquite analyses, Supporting Information, S8, yielded equivocal results). The RASP analysis (Supporting Information, S5) indicated that this character evolved at least five times independently within the order, once in Faboideae, three times within the rest of the legumes, and once in Polygalaceae. Within Faboideae, while the MRCA possessed non-enclosed reproductive organs (node 1019, B= 99.6%), polymorphic flowers (i.e, in clades with enclosed and non-enclosed reproductive organs, as in the *Xanthophyllum*) evolved after the separation of the Swartzieae clade (node 1013, B=92.2%, AB= 7.8%, 67.2 MYA), became dominant after the separation of the ADA clade (node 998, AB=69.4%, B= 1.4%, A=16.2%, 66.7 MYA). After that, while the percentage of polymorphic flowers decreased and flowers with enclosed reproductive organs became dominant gradually (nodes 998, 995, 993, 989, 983), enclosed reproductive organs were well established after the origin of the Genistoid s.l. clade of Cardoso *et al*. (2012a) (Node 979, A=78%, AB=22%, 62.5 MYA) (Figure 2). The molecular dating analysis indicated that this experimental phase lasted from 167.19 MYA (the origin of Faboideae) to 62.5 MYA (the origin of the Genistoid s.l. clade). Ater the evolution of the Genstoids, there were 13 reversals. These were: *Cadia* Forssk. (29.2 MYA), *Dicraeopetalum* (13.6 MYA), *Camoensia* Welw. ex Benth. & Hook. f. (25.9 MYA), *Ormosia* Jacks. nom. cons. (polymorphic flowers) (29.5 MYA), *Amphicarpeae* (26.1 MYA), *Baphiopsis* (19.7 MYA), *Airyantha* Brummitt (32.6 MYA), *Etaballia* Jacq. nom. cons. (10.8 MYA), *Inocarpus* J. R. Forst. & G. Forst. (11.3 MYA), *Riedeliella* (30.1 MYA), *Acosmium* (59 MYA), Amorpheae clade (15.2 MYA) and *Aldina* (62.9 MYA) (note that *Sophora*+*Calia* clade is outside the Genistoid s.l. clade in the analysis; however, this phylogenetic difference does not affect the result for the origin of the family).

Within the rest of the legumes, flowers with enclosed reproductive organs independently evolved three more times, namely in *Cercis* (39.3 MYA), *Pomaria* (25.5 MYA), and at the origin of *Tara*+*Coulteria* (13.7 MYA). On the other hand, the RASP results both with ML and BI starting trees, indicate that an experimental phase might also be possible for the family Polygalaceae. While the MRCA of the family possessed polymorphic-flowered populations (node 1262, AB=97%, 63.59 MYA), enclosed reproductive organs were probably well established in the Polygalaceae after the separation of the *Xanthophyllum* and *Moutabea* clades (node 1260, A= 93%), about 49.5 MYA; with one revearsal (i.e., *Atroxima*+*Carpolobia* with polymorphic flowers) (node 1256, AB=92%), about 20.1 MYA.

Third, for the three different petal/petal+sepal types, all of the analyses yielded ancestors with one type of petal for Fabales (89.3% and 99.9%), Fabaceae (76.2% and 96.4%) and Polygalaceae (82.5% and 95.1%) (Table 2, Figure 2). Conversely, for the tribe Polygaleae, the MRCA probably possessed three types of petal, as indicated (89.6% and 99.0%) by the results. For the Faboideae, while the results of the Mesquite analysis (Supporting Information, S9) yielded equivocal results, the BayesTraits (two types of petal, 83.6%) and RASP (one type of petal, 97.8%) results conflicted.

According to the RASP analysis (Supporting Information, S6), while the origin of the Faboideae did not possess three types of petal (i.e., two types of petals) (node 1019, B=97.8%), flowers with three-types of petal were well-established after the separation of the Swartzieae (node 1013, A=87.7%, AB=9%). However, until the branch leading to the Genistoid clade (node 979; A=91.4%, AB=8.5%) in which the three-types of petals were well-established again, the percentage of three-types of petals suddenly decreased, and the percentage of polymorphic flowers increased (nodes 993, 989 and 983). Interestingly, this time frame coincides with the K-T (Cretaceous-Tertiary) extinction period (from ca. 67-19 to 64.79 MYA). However, after the establishment of three-types of petals within the family, only 12 reversals and just two to only one type of petal are noteworthy. The evolution of non-papilionoid-flowered clades (i.e., Swartzieae, ADA clade, *Cladrastris* clade) is also noteworthy. Of note is that the timing of the evolution of three-types of petals within Faboideae coincides with the evolution of the enclosed reproductive organs.

Within the rest of the Fabaceae, while three-types of petals evolved at least 15 times, two types of petals evolved at least 25 times, independently. For the Polygalaceae, on the other hand, while the family possessed flowers lacking-three-types of petals at its origin, three-types of petals evolved at the origin of tribe Polygaleae+tribe Carpolobieae (node 1257, B=34%, A=27%, C=26%) and in *Xanthophyllum*, and were well established at the origin of tribe Polygaleae (node 1255, A=99%).

## DISCUSSION

Since the early-branching non-keeled Faboideae genera follow different developmental origins (i.e., homoplasy is present) (e.g., Pennington *et al*. 2000), instead of coding flowers as keeled or non-keeled, I divided the keel flower syndrome into three further characters, widely acknowledged in numerous studies as pivotal morphological traits of this flower type (e.g., McMahon and Hufford, 2005; Carvalho *et al*. 2023): the presence of enclosed reproductive organs, the presence of bilateral symmetry, and the perianth heteromorphism (Pennington *et al*. 2000; Tucker, 2002; Westerkamp, 1997; Cardoso *et al*. 2012a). While including some other characters of the keel flowers, such as petal aestivation (Tucker, 2002), organ initiation pattern (Pennington *et al*. 2000), and hypanthium (Pennington *et al*. 2000) are desirable; due to limited information on these characters in the literature these are not included in the current study. Nevertheless, the RASP analyses, in particular, yielded results bearing on floral evolution within the order Fabales.

### Floral symmetry

The symmetry analyses indicated inconclusive results regarding the origin of the order Fabales. It is important to note that this ambuigity may be a result of character coding, specifically the distinction between bilateral symmetry and radial/slightly bilateral symmetry. The results should be interpreted as a “radially or slightly bilaterally” symmetrical origin.

In this context, the detailed analyses of Bello *et al*. (2012) reported that bilateral symmetry likely evolved independently within legumes and the tribe Polygaleae. Reyes *et al*. (2016) proposed that the origin of Fabales flowers could be radial symmetrical, while legumes in general and Faboideae in particular most probably had an origin in bilateral symmetry. Similarly, Sauquet *et al*. (2017) reported a radial symmetrical ancestor for Fabales, although sampling in that study was low. Further testing is warranted to explore this aspect more comprehensively.

Similarly, the analyses produced equivocal results concerning the MRCA of Faboideae. Whilst the result of the Bayestraits MCMC ancestral trait reconstructions moderately suggest a radial or slightly bilateral symmetry (only 80.1%) for the origin of the Faboideae, the remaining analyses strongly suggest that bilateral symmetry evolved before the evolution of the subfamily, at the origin of the (Fabaceae+Polygalaceae) node.

Citerne *et al*. (2006) reported that even in radially symmetrical Faboideae flowers, such as those of *Cadia,* the initial developmental stages exhibit bilaterally symmetrical flowers, followed by the subsequent development of radial symmetry. This observation was further supported by several studies emphasizing that zygomorphic symmetry is expressed early in papilionoid flower development, with both keel and non-keeled flowers following the same basic pattern for symmetry expression (Tucker, 1997, 1999, 2002; Pennington *et al*. 2000; Lavin *et al*. 2005; LPWG, 2017) (also refer to Tucker, 2002; Cardoso *et al*. 2013 and Bento *et al*. 2021). Therefore, the radial or slightly bilateral symmetry results of the BayesTraits analyses for the ancestral flower type of subfamily Faboideae could be artefacts of the methods used, potentially arising from character state assignment (Anonymous Reviewer 2, pers. comm.), incomplete taxon sampling, and/or if the trait positively effects the diversification rates within the clade (Maddison *et al*. 2007; Leys and Hogendoorn 2008; Ron, 2008). It is plausible that the original flower form of the subfamily was bilaterally symmetrical (Endress, 1999) at least to some degree.

Conversely, most of the closest relatives of Faboideae, in the Caesalpinioideae, have radially/slightly bilaterally symmetrical flowers (about 100 genera out of 163 genera). It is possible that, the evolution of strongly bilaterally symmetrical flowers occured after the origin of Faboideae, as in the evolution of enclosed reproductive organs and petal heteromorphism (discussed below). Indeed, both the character reconstructions and recent ontogenetic studies (e.g., Table 2 of Bento *et al*. 2021) have shown that not only keel flowers, but also non-keeled flowers of Faboideae undergo remarkable changes in symmetry during corolla development, which indeed may suggest a family-wide ongoing experimental phase *sensu* Prenner and Klitgård (2008).

The RASP results also indicated that reversals (or evolutionary innovations with diverse mechanisms, Citerne *et al*. 2006; Sinjushin, 2019; Bento *et al*. 2021) from bilateral symmetry to radial symmetry were quite common, as also seen in other angiosperm families (Boatwright *et al*. 2008; Reyes *et al*. 2016); even within Faboideae (at least 13 times lost, and three times re-gained) and Polygalaceae (lost twice), but particularly within the remaining members of the Fabaceae. The results support the analyses of Bruneau *et al*. (2014) and Reyes *et al*. (2016) suggesting that reversals to radial symmetry are more common in Fabales and particularly in caesalpinioids, when compared to other angiosperm families and legume subfamilies. Bruneau *et al*. (2014) also concluded that these reversals may be easier for the caesalpinioid clades, particularly concerning reductions in organ number, organ initiation patterns, and symmetry, where flower specialization is not as extensive as in papilionoids (Pennington *et al*. 2000; Leite *et al*. 2015). Indeed, the authors postulated 11 symmetry changes, 27 stamen number changes and 15 corolla morphology changes just in Detarieae (i.e., subfamily Detarioideae); but this variability in Faboideae and the mimosoid component of Caesalpinioideae was less common. Interestingly, reversals to ancestral states for the two characters presence of enclosed reproductive organs and three types of petals were not as common as reversals back to radial symmetry.

### Enclosed reproductive organs

The possession of a flag petal (vexillum) and enclosed reproductive organs are arguably two of the most important characters for defining keel flowers, as previously suggested by several studies (e.g., Westerkamp and Weber, 1999; Carvalho *et al*. 2023). The character reconstruction of the presence of enclosed reproductive organs indicated that the origin of Fabales possessed non-enclosed reproductive organs; however, noteworthy is the fact that this character has evolved multiple times within the order.

Result from the RASP analyses indicate that the enclosed reproductive organs evolved at least five times independently within the order, once in Faboideae, three times within the rest of the legumes, and once in Polygalaceae. While the ancestral node of the subfamily Faboideae reconstructed to have non-enclosed reproductive organs, as all of the analyses strongly suggested (99.5-99.9%); polymorphic flowers (i.e., keeled and non-keeled flowers in the same clade(s), as in the case of *Xanthophyllum*and *Sophora*) evolved after the separation of the Swartzieae clade. These polymorphic populations became dominant after the separation of the ADA clade, while the protected reproductive organs did not become dominant until the branch leading to the crown Genisitioids. Therefore, contrary to recent views (e.g., Pennington *et al*. 2000; Lavin *et al*. 2001; Citerne *et al*. 2006; Cardoso *et al*. 2012a) the results of the present study support the earlier assumption (e.g., Arroyo, 1981; Polhill & Raven, 1981) that the enclosed reproductive organs evolved slightly later in the history of the subfamily, not at the origin. And after the first appearance of the enclosed reproductive organs in the family, the stabilization of the character spanned 4.7 million years (from 167.19 MYA, the origin of Faboideae to 162.5 MYA, the origin of the Genistoid s.l. clade). Surprisingly, the molecular dating results, incorporating several fossil calibrations suggest that this experimental phase (or instability syndrome *sensu* Sinjushin, 2021) with polymorphic-flowered populations corresponds to the K-T mass extinction event (see below). Here, it should be noted that while my analysis identified Swartzieae as the earliest diverging clade, similar to the plastid phylogenomic studies of Zhang *et al*. (2020) and Choi *et al*. (2022), a recent unpublished study by Cai *et al*., which utilized 1,456 low-copy nuclear loci, produced a different result, identifying the ADA clade as the earliest diverging group within Faboideae.

Thirteen reversals (between ca. 62.9 and 10.8 MYA) occurred later within the subfamily, possibly due to one or more of the following causes: pollinator inefficiency, pollinator limitation, pollinator shifts (Pennington *et al*. 2000; Klitgård *et al*. 2013; Breitkopf *et al*. 2015) or spatiotemporal fluctuations of pollinators (Fenster *et al*. 2004); ecological and spatio-temporal distributions (e.g., geographic ranges, relative abundance and flowering times), (Joffard *et al*. 2019) and physical environment which may influence the plant-pollinator interactions (Sargent and Ackerly, 2008); competition for ecological niches (i.e., adaptation to new habitats) or competition for limited sources in stressful conditions (Sargent, 2004; Pellissier *et al*. 2010; Althoff *et al*. 2014); and developmental and/or ecological factors (McMahon & Hufford, 2005). Interestingly, almost all non-keeled flowered lineages of Faboideae have a tropical distribution (Allen and Allen, 1981), such as wind pollinated *Ateleia*. (Janzen, 1989; Pennington *et al*., 2000); bird pollinated *Cadia*, *Myroxylon balsamum* L. (Harms), *Castanospermum australe* A. Cunn. & C. Fraser ex Hook. and tribe Angylocalyceae (Tucker, 1993, 1994; Pennington *et al*., 2000; Boatwright *et al*., 2008; Cardoso *et al*., 2012). It has been reported that bees are less dominant in wet environments (Pellissier *et al*. 2010) and the abundance of bees can vary significantly even within small changes in elevation (Conrad *et al*. 2021). If the evolution of non-keeled flowers within the tropics is driven by pollinator limitation or a similar factor, as suggested by several authors (Pennington *et al*. 2000; Klitgård *et al*. 2013), I would expect to observe a similar pattern for the tribe Carpolobieae, the tribe represents the only reversal to a non-enclosed floral form in Polygalaceae. However, it is important to note that the exact ancestral biogeographical region of this clade remains unknown and requires further testing (Aygören Uluer *et al*. 2022a reported an African origin for this clade).

In contrast, the evolution of keel flowers has been linked to pollinator specialization, as proposed by various researchers (Arroyo, 1981; Westerkamp, 1996, 1997). This is because with increasing floral complexity functional pollinators tend to become more restricted (Fenster *et al*. 2004). If this specialization occurred just before the K-T boundary (i.e., before the separation of Swartzioids, ca. 67.2 MYA), it is plausible that subfamily Papilionoideae passed through an experimental period with different flower types, or the evolution of enclosed reproductive organs could have increased the visits of new pollinators (Dell’Olivo & Kuhlemeier, 2013; Liu *et al*. 2019). However, this hypothesis requires further testing and investigation.

For the Polygalaceae, a second experimental phase lasted for 14.1 MY. While polymorphic-flowered lineages were dominant in the ancestral population(s) of the family (i.e., as in the extant genus *Xanthophyllum* with keel and non-keeled flowers) as the results suggested, the dominance shifted to enclosed reproductive organs later in the evolution of the family. This experimental phase not only lasted for a more extended period compared to the Faboideae (14.1 MY vs. 4.7 MY, respectively), but also occurred after the evolution of the enclosed reproductive organs in Faboideae (i.e., just after the K-T boundary, 63.6 to 49.5 MY). The duration of this experimental phase might have been influenced by the distinct floral bauplan of the keel flowered Polygalaceae or a possible mimicry scenario suggested by some studies (e.g., Tucker, 2002; Aygören Uluer *et al*. 2022a). For instance, similar to some Faboideae (e.g., *Taralea* Aubl., *Pterodon* Vogel and *Dipteryx* Schreb. with petaloid adaxial sepals in Dipterygaeae; Leite *et al*. 2014; Carvalho *et al*. 2023) sepals have a role in the composition of the keel flowers in Polygalaceae (Westerkamp, 1997; Westerkamp and Weber, 1999).

Is it possible for a keel flowered ancestor to have exposed reproductive organs or undifferentiated petals? If the keel plays a crucial role in the reproductive biology of these plants by facilitating efficient pollination—enhancing pollen transfer and ensuring the plant’s reproductive success through specific pollinators such as bees—along with ensuring the pollinator’s contact with reproductive organs during landing, then the evolution of both the keel and the enclosed reproductive organs become perhaps the most important criteria in defining a keel flower (Westerkamp, 1997). Here, while Endress (1996) suggested that, albeit rarely, some keel flowers can be non-enclosed, Tucker (2002) eloquently explained the definition of keel flowers using adjectives such as “barely, scarcely, imperfectly, strongly, and pseuodopapilionoid”. For example, a keel is formed by the curving inwards of the barely differentiated keel petals (Tucker, 2002), in *Poeppigia procera* (Poepp. ex Spreng.) C. Presl and by the reflexion of the upper petals, a pseudopapilionoid floral morphology is expressed (Tucker, 2002). Therefore, I believe that by definition (e.g., Arroyo, 1981; Polhill & Raven, 1981; Howell *et al*.1993; Endress, 1996; Westerkamp, 1996, 1997; Westerkamp & Weber, 1999; Pennington *et al*. 2000; Persson, 2001; Tucker, 2002; Westerkamp & Claßen-Bockhoff, 2007; Bento *et al*. 2021), having protected reproductive organs lies at the core of the definition of a keel flower.

### Heteromorphism in sterile organs

The third character is the presence of three or less different types of petals. One-type of petal at the origin of the Fabacea eis moderately to strongly supported (i.e., 76.2-96.4%), the origin of subfamily Faboideae had one or two different petal morphologies, and the origin of the Polygalaceae with only one type of petal. Moreover, the RASP analyses revealed that this character evolved at least 18 times within Fabales: once in Faboideae, 15 times within the rest of the legumes, and twice in Polygalaceae.

The presence of conflicting results among various ancestral state analysis methods is a well known issue (e.g., Kondraskov *et al*. 2015). Assuming that the findings are not a result of artefacts of the methods used, they suggest that the origin of Faboideae did not possess three-types of petals. Instead, this character appears to have evolved after the separation of the *Swartzieae* clade, and later became dominant, experienced regression around the K-T boundary (ca. 67.19 to 64.79 MYA) and re-emerged as the dominant trait around 62.5 MYA on the branch leading to the Genistoid clade. Notably, this timeframe (i.e., ca. 67.19 to 62,5 MYA) coincides with the first appearance of enclosed reproductive organs within Faboideae (Figure 2).

While petal heteromorphism into three types is the predominant form in Faboideae flowers, and a floral ancestor with three types of petals has been suggested in previous studies for the origin of the subfamily (e.g., Pennington *et al*. 2000), the results indicate that the origin of the subfamily probably possessed only one (RASP, 97.8%) or two types of petals (BayesTraits, 83.6%). In this context, Tucker (1997) proposed that petal size and shape differences do not emerge until the midstage of the organ development in papilionoid flowers. If the hierarchical hypothesis is correct (but see Bento *et al*. 2021), this may further suggest the presence of only one type of petal at the origin of the subfamily. However, it is important to note that even without differentiation in petal size and shape, variation in position may still be possible e.g., a standard-like structure formed during petal aestivation. This is evident in fossil flowers of *Barnebyanthus buchananensis* Crepet & Herendeen (Crepet and Herendeen, 1992), dated to ca. 56 MYA, *Cercis* with barely differentiated petals (Tucker, 2002), and with a slight shape difference as seen in the pseudo-papilionoid flowers of *Poeppigia* (Falcão and Mansano, 2021).

If the age of the subfamily is estimated to be 67.19 MYA (95% HPD 62.5–64.9), the MRCA of Faboideae could have possessed flowers with a caesalpinioid-like or early-diverging papilionoid like morphology. These flowers might have been zygomorphic, with only one type of petal (possibly including a keel-like petal that at least partly covered the reproductive organs or the standard having outermost aestivation) and exposed reproductive organs, as in many caesalpinioids (e.g., *Duparquetia* Baill., *Poeppigia procera*, *Peltophorum* pterocarpum (DC.) K. Heyne and *Bauhinia* L.). Crepet and Herendeen (1992) placed the *Barnebyanthus buchananensis* fossil in Sophoreae, a taxonomic group no longer in use in its traditional sense but accepted as transitional between Swartzieae and Faboideae. The *B. buchananensis* fossil flowers (Faboideae) dated as Late Paleocene to Early Eocene (ca. 56 MYA), (Crepet and Herendeen, 1992) are bilaterally symmetrical, with exposed stamens, and all petals alike except for the outermost standard (i.e., differentiation into a standard, but not wings and keel). This floral structure may be similar to the MRCA of Faboideae.

The case of *Poeppigia* is noteworthy. While the reproductive organs are enclosed in bud by barely differentiated petals, they become partly exposed in fully blooming flowers, yet somehow restrict access for the bees to the reproductive organs (Tucker, 2002; Falcão and Mansano, 2021), similar to *Zollernia* Maximil. & Nees (Faboideae), *Coulteria* Kunth. and *Tara* Molina (Caesalpinioideae). Indeed, the morphological similarity between the fossil *Barnebyanthus buchananensis a*nd *Cercis* flowers is striking.

In comparison, even though only moderately supported, the possibility of differentiation into two types of petals in the BayesTraits analyses should not be underestimated, especially because two types of petals are not uncommon in subfamilies of Fabaceae, being particularly prevalent within Caesalpinioideae (e.g. *Caesalpinia* L.). It is also observed in subfamilies Cercidoideae, Detarioideae, Dialioideae and Duparquetioideae (e.g*., Hymenaea* L. in subfamily Detarioideae, *Martiodendron* Gleason in subfamily Dialioideae, *Adenolobus* (Harv. ex Benth.) Torre & Hillc. and *Bauhinia* in subfamily Cercidoideae. Two petal types are also seen in *Castanospermum,* and *Errazurizia megacarpa* (S. Watson) I.M. Johnst in the subfamily Faboideae. Nevertheless, the character is not unique to Fabaceae and Polygalaceae; it occurs in other angiosperm families (Tucker. 2002; Prenner & Klitgård, 2008; Bruneau *et al*. 2014). For instance, both *Aconitum* L. (Ranunculaceae) and *Corydalis cava* (L.) Schweigg & Körte (Fumariaceae) have petaloid sepals and petals that are clearly differentiated into three types as in keel flowers (Proctor *et al*. 1996). Therefore, while it is evident that further analyses are needed, I believe that the ancestral floral form of Faboideae might have possessed one broad standard petal and four similar narrow petals (i.e., two types of petals) or at least a standard petal without any size or shape difference. This standard petal, however, most likely occupied a specific position relative to the other petals, making the flower at least moderately bilaterally symmetrical (e.g., *Myroxylon* L. *Angylocalyx* Taub. *Castanospermum* and *Camoensia*) (note that the analyses yielded mostly equivocal results for this character).

Numerous angiosperm studies have demonsrated that the K-T event resulted in extinctions not only within plants and animals but also within plant-polllinator relationships, often followed by subsequent radiations (e.g., Rehan *et al*. 2013 and references therein). For instance, Rehan *et al*. (2013) specifically highlighted extensive extinction events within the species numbers of the bee subfamily Xylocopinae Latraille as a direct consequence of the K-T extinction event. Apidae Latreille (Xylocopinae) is one of the two bee families documented to pollinate Fabales keel flowers (the other family being Megachilidae), particularly the genera *Apis* Linnaeus, *Anthophora* Latreille, *Eucera* Scopoli, *Xylocopa* Latreille, *Bombus* Latreille and *Centris* Fabricius (Aygören Uluer, 2021). In this context, the results indicate a regression of petal differentiation and an experimental phase with polymorphic flowered populations around the K-T boundary which occurred ca. 66 MYA (Wilf *et al*. 2003; Rehan *et al*. 2013). If the main pollinators of keel flowers became extinct during this timeframe, it is possible that the disturbed plant-pollinator relationships led to the acquisition of more generalist pollinators and dominance of non-enclosed reproductive organs (i.e., more open flowers) for a relatively short period. However, the assumption that non-keel flowers are pollinated by a broader array of pollinators has not been extensively tested, except in a few local studies. For example, while *Amorpha canescens* was visited by several different insects, a nearby *Baptisia* Vent. species with keel flowers was visited by only a single bee species (Robertson, 1890, McMahon and Hufford, 2005). A recent study reported opposite results, indicating keel flowers of *Discolobium pulchellum* Benth. and non-keeled flowers of *Riedeliella graciliflora* Harms., both pollinated by bees; however, the behaviour of the bees differed between the two species (Bérgamo, 2017; but refer to Bento *et al*. 2021 for further details on this behavioural difference). Therefore, field observations of pollinators of keel and non-keel flowers of papilionoids, as well as reconstructions of the possible ancestral pollinators, are needed to test these plant-insect relationships. Also, as McMahon and Hufford (2005) suggested, other genetic and ecological factors might have accompanied the changes in pollinator.

Secondly, the results indicate that both the origin of the flowers of legumes and Polygalaceae possessed one type of petal. In the case of legumes, both two and three types of petals evolved multiple times in individual lineages (e.g., subfamilies Caesalpinioideae and Faboideae), or within specific species. For the family Polygalaceae, as anticipated, petal heteromoprhism evolved at least twice, in *Xanthophyllum* and at the origin of Tribe Polygaleae+ Tribe Carpolobieae. The results suggest that the morphological traits attributed to the origin of a taxonomic group may have evolved multiple times within that clade and/or emerged not at the clade’s origin.

### Age of Fabales

It should be noted that the age estimations of the subfamily Faboideae vary in the literature, ranging from 45 to 58.6 MY (Lavin *et al*. 2005, Bruneau *et al*. 2008, Koenen *et al*. 2021; Calvalho *et al*. 2023). While almost all these studies included several fossils (e.g., up to 20), the ingroup sampling was sufficient only in Lavin *et al*. (2005). In contrast, the results of the current study suggest an older age for the subfamily, specifically 67.19 MY (95% HPD 62.5–64.9), employing 30 fossil calibration points and a very large ingroup sampling.

As Koenen *et al*. (2013) proposed, a significant number of fossils with the same age (45-46 million years) might have negatively influenced the analysis, considering nine fossils in the present study. The importance of taxon sampling in molecular dating analyses has been demonstrated in various studies (e.g., Linder *et al*. 2005; Heath *et al*. 2008). To address this, the present study incorporated the largest outgroup sampling to date for a more balanced tree (Smith, 1994). However, the impact of fossils, both ingroup and outgroup sampling on Fabales molecular clock analysis remains uncertain (Koenen *et al*. 2013). Further studies with well-sampled taxa (and data), and robust fossil calibration points are needed.

### Evolution of keel flowers within Fabales

The number of non-keeled flowers in the early-branching Faboideae does not guarantee a non-keeled origin. Non-keeled genera are scattered throughout the Faboideae phylogenetic tree and the early-branching non-keeled genera follow different developmental origins (i.e., homoplasy) (e.g., Pennington *et al*. 2000; Cardoso *et al*. 2013). For instance, with respect to petal number, a non-keeled taxon can have undergone petal abortion (i.e., no petal primordia initiated), examples include: *Ateleia*, *Bobgunnia* J. H. Kirkbr. & Wiersema, *Swartzia* Schreb., *Aphanocalyx* Oliv. (Tucker, 1990, 2000, 2003b) and *Parryella* (McMahon & Hufford, 2005), or petals can be initiated but later suppressed (i.e., they are rudimentary), such as in *Amburana cearensis* (Allemão) A.C. Sm. (Leite *et al*. 2015). Additionally, both the ADA and Swartzioid clades include many different floral arrangements (e.g., Tucker, 1990, 1994; Leite *et al*. 2015), including an entire calyx and numerous stamens (e.g., *Cordyla*), reduced corolla (e.g., *Swartzia*), unidirectional organ initiation (e.g., *Ateleia*) and unusual ring meristem (e.g., *Mildbraediodendron*) (Tucker, 1992, 2000, 2003b; Sinjushin, 2018). It is also worth noting that in Polygalaceae, either five sepals and petals are initiated after which lateral petals are suppressed (Prenner, 2004) or lateral petals are completely lost (Bello *et al*.2010). However, these apomorphies are rare (Leite *et al*. 2014, 2015), and the number of organs is determined at the very early ontogenetic stage of keel flowers, while organ suppression takes place at the mid-stage (Tucker, 1997). Furthermore, these diverse ontogenetic pathways and morphologies of non-keeled flowers do not essentially point to a keel flowered ancestor; indeed, they could easily reflect the plasticity of early-branching Faboideae (Klitgård *et al*. 2013) or differing evolutionary pathway (Prenner & Klitgård, 2008). Similar hypotheses have been made for both early-branching legumes (Prenner & Klitgård, 2008) and early-branching Faboideae, highlighting the intricate evolutionary trajectories within these groups (Leite *et al*. 2015; Prenner *et al*. 2015; Ramos *et al*. 2016).

Supporting the hypothesis that subfamily Faboideae probably evolved from caesalpinioids (Leite *et al*. 2015), is that a flower bauplan featuring bilateral symmetry, non-enclosed reproductive organs, and pentamerous undifferentiated petals is a combined morphology common in caesalpinioids (e.g., *Caesalpinia*, *Senna* Mill. and *Cassia* L.) as well as in many early-branching Faboideae (e.g., *Zollernia*, *Castanospermum* and *Xanthocercis* Baill.). Flowers with enclosed reproductive organs and one or two different petal types might have evolved at a very early stage in the evolution of the subfamily, contributing to the significant floral diversity observed in early-diverging Faboideae lineages. If the origin of enclosed reproductive organs occurred within the time frame of 167.19- 62.5 MYA, this phase could have lasted for up to ∼4.69 million years. Indeed, the mass extinctions during the K-T boundary and the subsequent recovery period could have created new habitats for diversification, not only for Faboideae but for all Fabaceae (Renner & Schaefer, 2010; Koenen *et al*. 2013, 2021). It is possible that Faboideae could have used niche broadening during this period (Renner & Schaefer, 2010) taking advantage of its floral plasticity (Tucker, 2003b; Ojeda *et al*. 2019) and the significant capacity for flower evolvability within the early-branching legume lineages. These lineages exhibit unusual ontogenetic pathways, including organ reduction, organ formation followed by later suppression, or even no organ initiation (Klitgård *et al*. 2013; Leite *et al*. 2015). The variable floral morphologies observed today could be remnants of the events that occurred on the early branches of Papilionoid origin, with subsequent separations into early-diverging clades and potential later extinctions.

The evolution of enclosed reproductive organs extends beyond the subfamily Faboideae and tribe Polygaleae, occurring three times within other legume lineages. Although the convergent evolution theory has been proposed for these independent evolutions (e.g., Westerkamp, 1997; Westerkamp and Weber, 1999), the possibility of parallel evolution should not be disregarded (Vasconcelos *et al*. 2017), particularly with regard to enclosed organs and petal/sepal heteromorphism in different families of the order. Whether convergent or parallel, the multiple evolution of this complex flower, not only within Fabales but also in various other angiosperm families may explain the co-evolutionary dynamics between keel flowers and their Hymenopteran pollinators, as well as the underlying genetic factors (Baguette *et al*. 2020).

The results of the current study align with other recent studies, corroborating findings such as those of Bruneau *et al*. (2014), which suggested that symmetry reversals are more common and likely to occur easily in caesalpinioids (compared to papilionoids), which exhibit a more uniform and specialized floral development pattern, while mimosoids display no symmetry changes. Second, in line with McMahon and Hufford (2005) who proposed a keel flowered origin for the Amorphoid clade, the results indicate that the Amorphoid clade’s origin is likely to have been bilaterally symmetrical (96.0%,), with enclosed reproductive organs (77.9% or 21.8% polymorphic), and three different petal types (91.3% or 8.9% polymorphic). Non-enclosed reproductive organs evolved twice in the clade (once in *Dalea* L. and once in the amorphoids); undifferentiated petals and radial symmetry evolved once in amorphoids, with a reversal to zygomorphy in *Amorpha*. Third, the floral evolution and homoplasy within the Vataireoid clade, were also remarkable (note that this clade was not monophyletic in the analyses), as De Olivera *et al*. (2023) reported that at least 35 evolution events happened independently in pollen traits. In the analysis, the members of this Vataireoid clade also exhibited remarkable diversity in the evolution of enclosed reproductive organs, petal heteromorphism and stamen connectivity. Fourth, Carvalho *et al*. (2023) also reported a radially symmetrical and non-keeled ancestor for the ADA+Swartzieae clades which contain some unique floral morphologies. The results match the hypothesis that the origin of these two early diverging clades possessed a non-papilionoid but bilaterally symmetrical ancestor. Last, but not least, a recent study (Cai *et al*. unpublished) proposed a non-keeled origin for Faboideae, with several independent origins and losses influenced by phylogeny. The authors indicated that the first keel flowers emerged around 59 MYA, just three million years before the *Barnebyanthus buchananensis*fossil. As the study suggests, the time difference between my findings and theirs regarding the evolution of keel flowers within the subfamily may be attributed to differences in taxon sampling and the calibration points used.

However, it is possible that the results of the analyses might have been influenced by the character coding and/or potential extinctions that have not been detected during the early evolution of the clades. While detecting such extinctions is challenging, future studies could include documented homology assessments to address this concern. However, considering that changes in the number of whorls (e.g., petals, stamens) are rare among Papilionoideae (e.g., rudimenteary petals, loss of petals, no petal initiation, Leite *et al*. 2014, 2015), and the number of organs is determined at the very early ontogenetic stage of keel flowers, while organ suppression takes place at the mid-stage (Tucker, 1997), I believe that the impact of homology on results should be minimal. Nevertheless, future studies should focus on the gaps in the literature and conduct ancestral character analyses based on homology assessments.

## FUTURE DIRECTIONS

Unfortunately, ontogenetic studies and overall understanding of morphology of the early branching non-keeled genera of Faboideae, and especially South American taxa, are limited (Prenner *et al*. 2015). The same applies to most Polygalaceae genera (except the most speciose genus *Polygala*). Information on other angiosperm clades with keel flowers is also very limited. I believe that a complete species level-sampling should be the goal of future studies.

Further research is also required to better understand the pollinators of keel flowers of different angiosperm clades, as well as the pollination biology of non-keeled genera of Fabaceae and Polygalaceae, A recent study by Alemán *et al*. (2022) indicated that different species pollinated by the same pollinators develop similar qualitative and quantitative morphological characaters. This should be tested in detail in early branching Papilionoideae and Polygalaceae

To explore diversification-trait associations, the origins of keel flowers have been treated as independent evolutionary events, but the results of the current study highlight multiple gains and losses of the main characters of keel flowers in what can be considered as “experimental” phases in floral morphology, before keel flowered species became numerous. Although the ecological implications of such experimental phases have not been explored in depth, evolutionary-developmental interpretations have been presented. A predisposition to evolve keel flowers in those lineages suggests clustered homoplasy (i.e., the aggregated morphological evolution of three different petal types, enclosed reproductive organs, and a deep corollatube) and the possible occurrence of cryptic precursors (i.e., evolution of a trait only after the gain of an intermediate trait), as articulated by Donoghue (2005), Donoghue & Sanderson (2015) and Marazzi *et al*. (2012). Both clustered homoplasy and cryptic precursor scenarios seem likely during the evolution of keel flowers in Fabaceae, and the analyses suggest different origins for some important morphological traits of the keel flowers.

## ACKNOWLEDGMENTS

The author is grateful to the Republic of Turkey Ministry of National Education for funding. I am grateful to Dr. Colin Hughes (University of Basel), Scott Armbruster (University of Portsmouth), associate editors and the anonymous reviewers for their constructive suggestions.

## SUPPORTING INFORMATION

Supporting Information S1, input file for the enclosed reproductive organs analysis

Supporting Information S2, input file for the petal heteromorphism analysis

Supporting Information S3, input file for the symmetry analysis

Supporting Information S4, RASP results for the symmetry analysis

Supporting Information S5, RASP results for the enclosed reproductive organs analysis

Supporting Information S6, RASP results for the petal heteromorphism analysis

Supporting Information S7, Mesquite results for the symmetry analysis

Supporting Information S8, Mesquite results for the three distinct petal/petal+sepal types analysis

Supporting Information S9, Mesquite results for the enclosed reproductive organs analysis

